# SURFS and AlphaFold Reveal Ribosome Footprint Shift Caused by EF-Tu D81 Mutation

**DOI:** 10.1101/2025.06.05.657321

**Authors:** Yi Zeng, Jordan Johnson, Shoujun Xu, Yuhong Wang

**Affiliations:** Department of Chemistry, University of Houston, Houston, TX 77204, USA; Department of Biology and Biochemistry, University of Houston, Houston, TX 77204, USA

## Abstract

Protein synthesis relies on accurate mRNA decoding by tRNA, a process guided by EF-Tu. We investigated how mutations at a conserved residue, aspartate 81, affect EF-Tu function using GTPase assays, AlphaFold modeling, and quantum-sensing-based super-resolution force spectroscopy (SURFS). All D81 variants retained GTPase activity but impaired tRNA release, revealed by sub-nucleotide ribosome footprinting. AlphaFold3 modeling suggests that D81 mutations disrupt magnesium coordination and interaction with the sarcin–ricin loop in the GTP-bound state. AlphaFold2-based sequence–structure analysis indicates that D81 anchors coevolutionary constraints, and its mutation enables cryptic structural variation. These results show how a single conserved residue links catalytic coordination, allosteric communication, and evolutionary constraint, offering mechanistic insight into translation fidelity and demonstrating the utility of an unconventional force spectroscopy in probing ribosome dynamics.

## Introduction

Protein synthesis relies on accurate codon–anticodon recognition between mRNA and tRNA, facilitated by the translational factor EF-Tu in complex with aminoacyl-tRNA and GTP (ternary complex).^1^ Misfunction in EF-Tu are linked to neurodegenerative diseases, and also make it a target for antibiotics.^2,3^ Cognate tRNAs accelerate GTP hydrolysis, enhancing decoding fidelity through kinetic control.^4,5^ This process is regulated via allosteric coordination between two major ribosome-ternary complex interactions: the GTPase activation center on the 50S subunit and the decoding center on the 30S subunit.^6,7^ Upon arrival at the ribosome, the anticodon stem-loop of the tRNA pairs with the mRNA codon, forming a minihelix, which is recognized by three 16S rRNA nucleotides – A1492, A1493 and G530.^8,9^ Meanwhile, the GTP-binding pocket of EF-Tu docks onto the sarcin–ricin loop (SRL) of the 50S subunit, positioning it for GTP hydrolysis.^10^ These transient interactions generate structural tension within the tRNA scaffold, inducing a kink in the anticodon stem-loop and a widening between the D- and T-loops.^11,12^ Upon GTP hydrolysis, EF-Tu undergoes a conformational shift, loosening its grip on the tRNA and allowing the release of torsional strain. This enables full accommodation of the tRNA, including its aminoacylated end, into the A site, preparing it for peptide bond formation. Meanwhile, EF-Tu dissociates from the ribosome, clearing the binding site for the next elongation factor.^5^

The GTP-binding pocket of EF-Tu contains five key motifs essential for its function: the P-loop, which interacts with the β- and γ-phosphates of GTP/GDP; switches I and II, which intercalate with the γ-phosphate; and motifs G4 and G5, which contribute to nucleotide specificity and binding stability (Figure 1A).^13^ These switches are highly flexible and serve as molecular sensors for GTP hydrolysis, undergoing significant conformational changes during EF-Tu activation. A critical residue Aspartate 81 (D81) in switch II, plays a key role in stabilizing the γ-phosphate by coordinating a Mg²⁺ ion (Figure 1B).^14^ Despite its conservation, D81 remains relatively underexplored, due to its mutation influencing EF-Tu stability.^15^ In this report, we generated three D81 mutants: D81A, D81F, and D81K, to investigate how changes in side chain size and charge affect EF-Tu activity. Despite preserving GTPase activity, all three mutations led to restricted ribosomal footprint motion on the mRNA in tRNA accommodation, as detected by SURFS spectroscopy with sub-nucleotide resolution. AlphaFold3 simulations suggest that the disruption arises from unstable Mg²⁺ coordination in switch II, where D81 normally stabilizes the Mg²⁺ ion and supports nucleotide binding. This instability may also change EF-Tu’s interaction with the sarcin ricin loop in the GTP bound form, but not in the GDP bound form, potentially blocking effective allosteric coupling to the decoding center. Sequence coupling to conformation was explored using AlphaFold2. Although the D81 mutation does not alter its apparent conservation in the multiple sequence alignment (MSA),^16^ it perturbs the surrounding coevolutionary network, allowing cryptic sequence diversity to emerge that was previously masked by entrenchment. This suggests a new application of AlphaFold: targeting conserved residues that anchor coevolution networks can uncover deeper structural variation not captured by sequence conservation alone.

**Figure 1.**
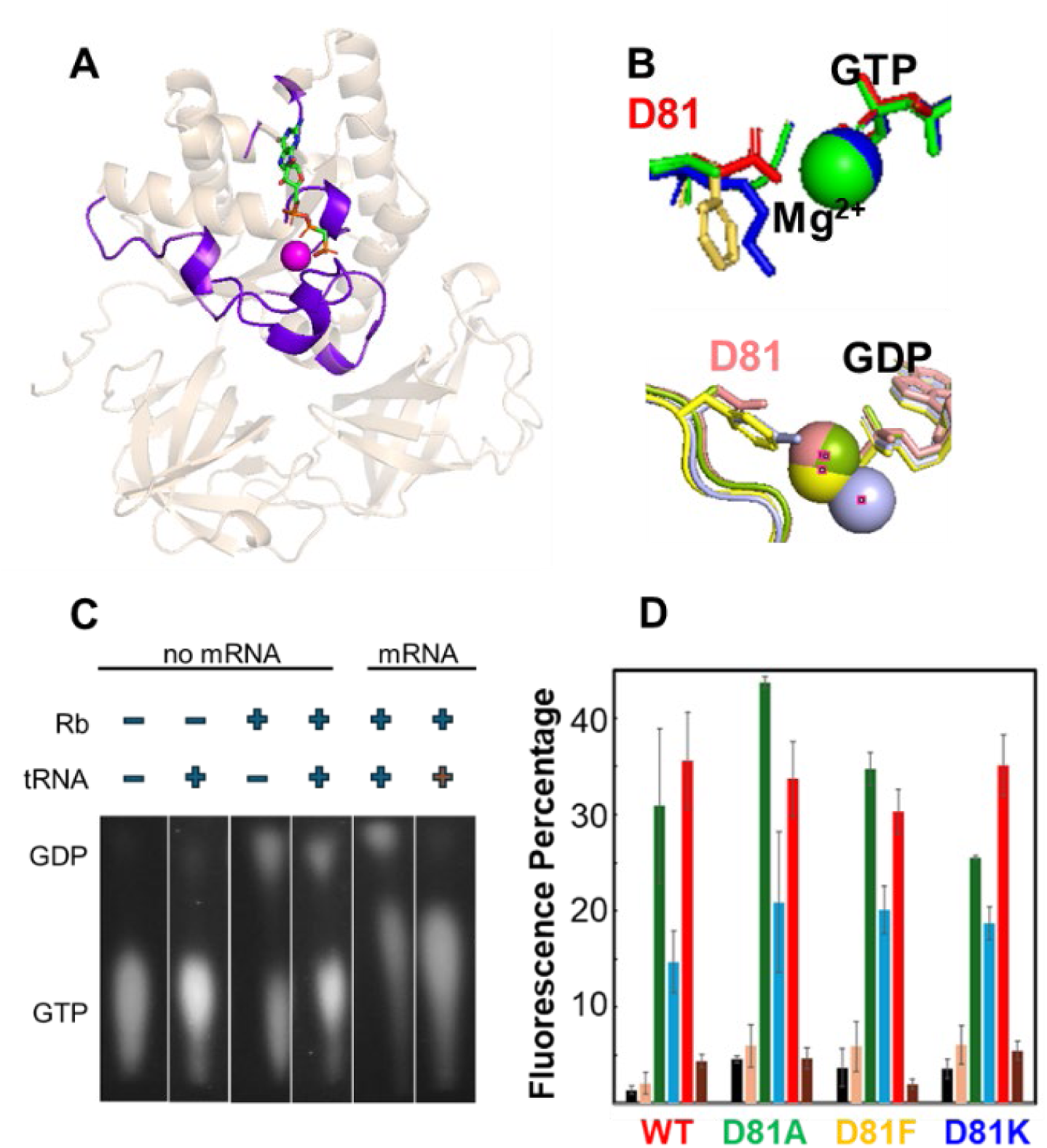
GTP hydrolysis center of EF-Tu. (A) The five conserved motifs involved in nucleotide coordination within the GTP binding pocket. (B) AlphaFold3 simulations showing the conformations of D81 and its mutants in GTP- and GDP-bound states. (C) Representative TLC images of mant-GTP hydrolysis by EF-Tu. (D) Bar plot summarizing GTPase activity across different conditions.

## Results

### GTP hydrolysis is not affected in mutants

We quantified GTP hydrolysis by measuring mant-GDP levels using fluorescence TLC, where higher percentage of mant-GDP indicates increased hydrolysis. Six conditions were tested (Figure 1): EF-Tu alone or with tRNA, with or without ribosome, and in the presence of mRNA with cognate or near-cognate tRNA. Without ribosome, GTP hydrolysis was minimal (with somewhat higher activities in mutants), consistent with EF-Tu’s low intrinsic GTPase activity. Ribosome presence significantly increased hydrolysis, confirming its role as a GTPase-activating factor (GAP).^10^ Aminoacyl-tRNA slightly suppressed hydrolysis in the absence of mRNA. Conversely, with mRNA, cognate ternary complex enhanced GTPase activity, while near-cognate tRNA suppressed it, demonstrating that EF-Tu’s GTP hydrolysis is coupled with proofreading. The D81A, D81K, and D81F mutants exhibited GTP hydrolysis patterns comparable to wild-type EF-Tu, indicating that D81 does not significantly affect the core mechanism of GTP hydrolysis during cognate versus near-cognate tRNA recognition.

### Ribosome footprints on mRNA are different for mutants

During decoding, the 30S subunit undergoes head swiveling and domain closure upon cognate tRNA recognition, altering the ribosome’s physical footprint on the mRNA.^17,18^ This subtle position change on the mRNA was precisely mapped using super-resolution force spectroscopy (SURFS), a technique developed by our groups to directly measure translocation fidelity in a single-turnover reaction—an event that traditional bulk assays cannot resolve.^19–21^ As shown in Figure 2, Post-translocation ribosome complexes were probed at the 3’ and 5’ mRNA ends using DNA probes that formed 14 base-pair (bp) duplexes at the 3’ end and 15 bp duplexes at the 5’ end. These probes enabled precise tracking of ribosomal footprint shifts during tRNA accommodation, which converts the Post-translocation complex into a Pre-translocation complex. The conformational change of the 30S subunit following tRNA binding to the A-site alters the availability of mRNA nucleotides for base-pairing with the DNA probes, leading to measurable shifts in dissociation forces. At the 3’ end, where tRNA accommodation occurs, the mRNA-DNA duplex exhibited an 8.8 pN force decrease in WT EF-Tu, reflecting efficient tRNA delivery. This shift, less than a single nucleotide displacement, highlights the high resolution of the SURFS method.^20^ The force decrease was 7.4 pN in D81F, 4.6 pN in D81A, and 3.1 pN in D81K, suggesting that these mutations induce subtle but distinct conformational changes in the ribosome. At the 5’ end, an increase in force was observed, measuring 9.0 pN in WT, 6.6 pN in D81F, 5.1 pN in D81A, and 2.8 pN in D81K. This force increase correlates with 30S head swiveling toward the 3’ direction after cognate tRNA recognition. The consistent force increase at this site indicates that the ribosome-covered mRNA region remains structurally stable, with no detectable expansion or compression during tRNA accommodation. If the ribosome’s footprint on the mRNA expanded or contracted, the force shifts at the 3’ and 5’ ends would differ, rather than showing a coordinated change, a pattern we have observed before.^22^

**Figure 2.**
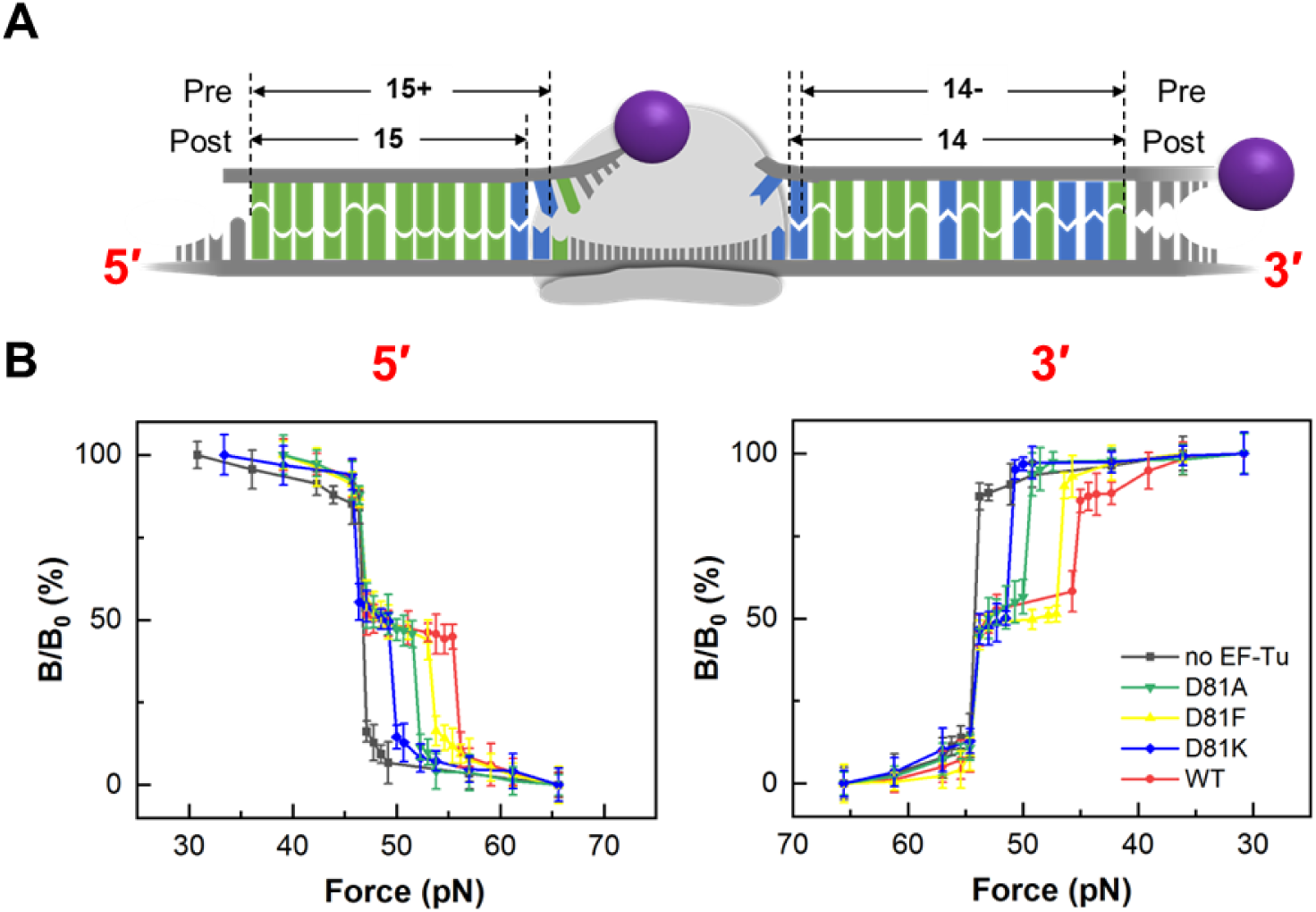
Probing EF-Tu function using SURFS from both the 5′- and 3′-ends (A) Probing scheme. Magnetic bead–labeled DNA probes hybridize to the exposed 5′ and 3′ ends of mRNA, and dissociation force reports ribosome position. (B) Force spectra of wild-type (WT) and mutant EF-Tu. The initial ribosome position was determined in the absence of EF-Tu (black traces), serving as a reference for EF-Tu–dependent shifts. Color code: WT-red, D81A-green; D81F-yellow; D81K-blue.

The progressive reduction in force shifts from WT to D81K at both ends suggests that D81 mutations disrupt EF-Tu’s ability to coordinate tRNA accommodation and ribosome movement, leading to impaired allosteric signaling between EF-Tu, the ribosome, and tRNA. In D81K, the weakest force changes at both sites suggest a less pronounced head swivel, potentially affecting proper tRNA accommodation, despite no detectable effect on their GTPase activities.

These results show that GTP hydrolysis alone does not account for the mutant effects. Instead, D81 mutations disrupt allosteric signal transmission between the GTP-binding pocket and the decoding center, impairing tRNA accommodation. D81K, which carries a charge opposite to that of WT’s aspartate residue, showed the weakest force shift, likely due to electrostatic disruption at the Mg²⁺ coordination site.

### AlphaFold3 simulations reveal mechanistic insights into EF-Tu mutant effects

Two key conformational changes underlie EF-Tu function (Figure S1). First, a global rearrangement between domain I and domains II–III (D-open vs. D-close): the closed state has high affinity for aminoacyl-tRNA, whereas the open state has reduced affinity.^23^ Second, a local shift in the switch I loop (SW-open vs. SW-close) facilitates Pi release after GTP hydrolysis.^24^ D81 at the adjacent switch II region coordinates a Mg²⁺ ion essential for GTP binding. While wild-type EF-Tu structures are well-characterized, the structural effects of mutations remain poorly defined.^25^

To investigate this, we folded EF-Tu–GTP and EF-Tu–GDP complexes with AlphaFold3.^26^ All models showed high overall confidence (Table S1). Tu–nucleotide interactions remained strong (iPTM 0.97–0.98), but Mg²⁺ coordination was weaker and more variable in mutants. D81K showed the most disrupted Mg²⁺ binding, with lower Tu–Mg²⁺ iPTM scores and higher PAE values (up to 3.7 Å). Nucleotide–Mg²⁺ geometry was also less stable in mutants. WT outperformed all mutants across structural metrics.

Inclusion of GTP, GDP, and Mg²⁺ alone predicted only the D-open conformation but not the expected D-close state of GTP-bound EF-Tu, although D-open state was also observed experimentally.^14,24,27^ To resolve this, we added two RNA elements—the sarcin-ricin loop (SRL, 30 nt) and a 22-nt RNA segment occupying the RNA-binding groove between domains II and III. These additions enabled AlphaFold3 to recover the D-close conformation across all variants (Figure 3A). Despite this, Mg²⁺ coordination remained disrupted in mutants (PAE: 3.65–4.98 Å, Table S2). More strikingly, SRL engagement in GTP-bound mutants was impaired (PAE: 6–14 Å), while GDP-bound forms retained proper SRL docking (Figure 3B). This indicates a mutation- and GTP-specific disruption of the interaction with ribosome GTPase activation center.

**Figure 3.**
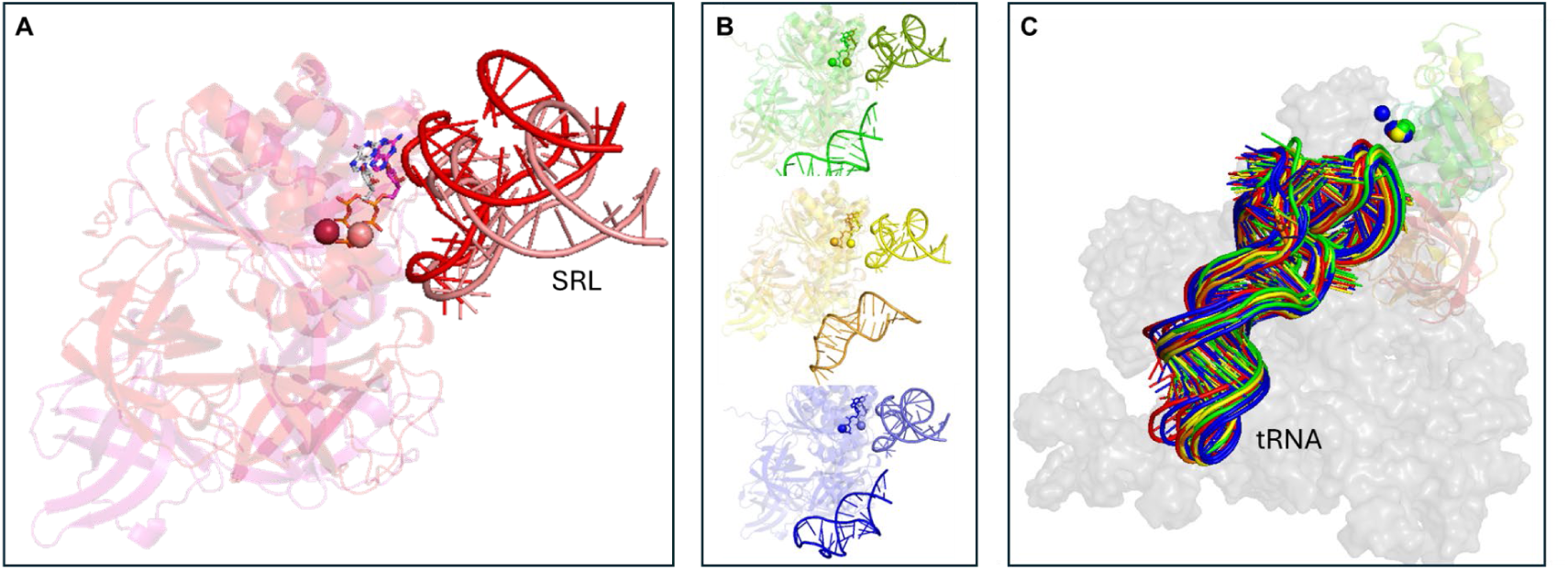
AlphaFold3 reproduces experimentally observed EF-Tu conformations aligned in domain I. (A) Overlay of closed (dark red) and open (light red) conformations of WT EF-Tu bound to GTP or GDP, Mg²⁺, 30-nt SRL, and 22-nt RNA (not shown for clarity). (B) Although both closed (dark) and open (light) forms are reproduced for the D81 mutants, the interaction between the SRL and the GTP-binding pocket is altered in the GTP-bound forms but remains unchanged in the GDP-bound forms. From top to bottom: D81A, D81F, and D81K. (C) Folding of EF-Tu in complex with partial 16S rRNA (nucleotides 5–650), tRNA (from PDB 5AFI), GTP, and Mg²⁺. All five AlphaFold3-predicted tRNA models are overlaid. All EF-Tus adapt close conformation.

We further modeled GTP-bound EF-Tu variants with 16S rRNA (residues 5–650) and tRNA (Figure 3C). The models aligned well with the cryo-EM structure (5afi).^28^ As before, Mg²⁺ coordination remained less stable in mutants (Table S3). Noticeably, one of five models in D81K showed a pronounced Mg²⁺ divergence. On the other hand, no major differences were observed in tRNA positioning, which may reflect a limitation of the model due to the absence of the full ribosomal context.

In summary, D81 mutations consistently weaken Mg²⁺ coordination while preserving Tu– nucleotide interactions. They also impair SRL engagement specifically in the GTP state, correlating with elevated auto-GTPase activity (Figure 1D). The displaced Mg²⁺ in D81K (Figures 1B and 3C) likely explains its largest footprint divergence from WT in SURFS data.

### Structure clustering of apo-WT and mutant proteins with AlphaFold2

To further investigate the structural and evolutionary basis of D81 conservation, we performed AlphaFold2-based structure clustering of EF-Tu in its apo form (without ligands or RNA).^29–31^ This approach isolates the protein’s intrinsic structural preferences, allowing us to identify sequence-encoded conformational states potentially critical for function (Figures 4A-D). Following multiple structure predictions, we developed an automated clustering algorithm to classify conformations objectively—an improvement over our earlier approach, which relied on visual inspection.^32^ First, predicted models with pLDDT scores above 70 were pooled and annotated by source; for example, "WT_090" refers to the wild-type EF-Tu structure from MSA cluster 090. Second, we performed partial alignment using domains II and III, which are largely invariant across all models. This alignment allowed us to isolate conformational variability in domain I—particularly its terminal helix (residues 183–199), which shows the most pronounced orientation shifts and serves as a key marker distinguishing D-open and D-close forms linked to tRNA delivery.^24,33,34^ Third, RMSD was calculated specifically within this region to maximize resolution of conformational differences. Figure S2A shows the overlay of all structures aligned on domains II and III, showing the divergence in domain I position. Figure S2B presents the RMSD matrix of range 183-199 (last helix of domain I), and its hierarchical clustering. This analysis resolved six distinct structural clusters. Among them, con3 in magenta was the most populated, followed by con2 in pink, con1 in brown, and con6 (blue). Clusters con4 and con5 contained only single representatives, D81F_090 and WT_035 respectively. Figure S2C presents a cleaner summary of all structural clusters, showing one representative structure from each protein variant. Figure S3 sorted the conformations based on SW-open and SW-close. It showed that except D81F_090 and con6 (blue) exhibited correlated D-open to SW-open form, all the other conformations exhibited uncoupled change in these two sites. All scripts and instructions for this structure-based clustering workflow and following phylogenetic analysis are available on our GitHub repository: https://github.com/ywang6000/EF-Tu-structure-clustering.

**Figure 4.**
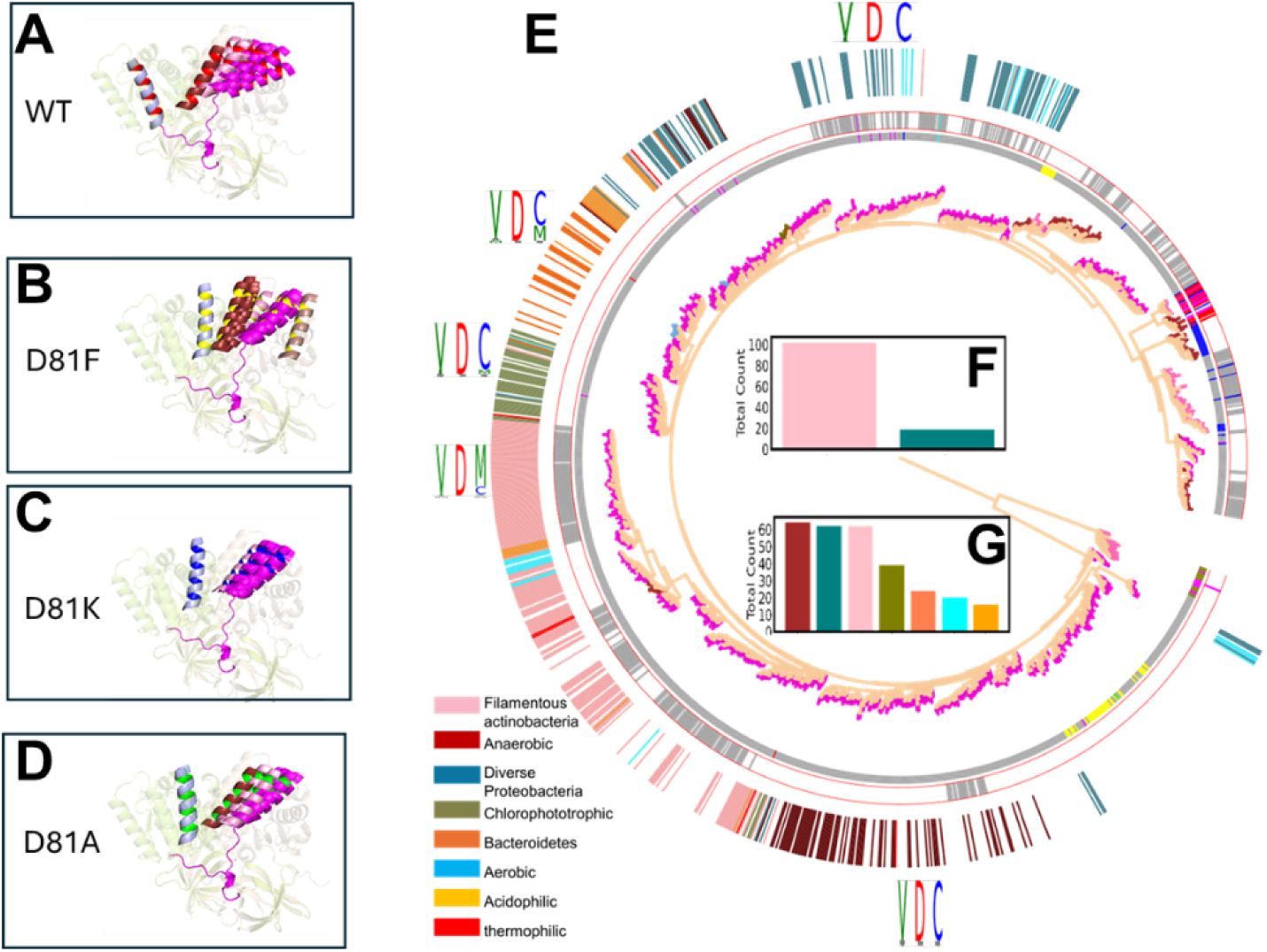
(A–D) High-confidence predicted structures using WT and D81 mutant sequences as queries. The transparent EF-Tu scaffolds are PDB 1EFC (GDP bound) and 1EFT (GTP bound).^14,36^ (E) Phylogenetic tree constructed after combining, de-duplicating, and labeling sequences. The inner two rings around the tree indicate whether each sequence is associated with D81K or WT, respectively. The outer ring represents species identity, with color codes shown in the plot. The three-residue LOGO plots (positions 80–82) show that position 81 remains conserved as aspartate. (F) Species analysis for sequences whose IDs contain “WT.” (G) Species analysis for sequences whose IDs contain “D81K” but not “WT.” Complete code is available at GitHub.com.

### AlphaFold2-guided phylogenetic analysis

We performed a structure-informed phylogenetic analysis by pooling only the sequences that yielded the high-confidence structures (Figures 4A-D). Protein identity, structure cluster, and query source are encoded in composite ID labels. For example, “EF-Tu-1_D81A_D81K_WT_A0A543IP01” indicates that UniProt ID A0A543IP01 is an EF-Tu protein adopting conformation 1, and it appears in the MSAs of three query conditions: D81A, D81K, and WT. The resulting unique sequence set was used to construct a maximum likelihood phylogenetic tree (Figure 4E).^35^ The tree resolved into two major branches (middle point root): one representing mostly EF-Tu (1197 leaves), predominantly associated with magenta conformations, and the other representing mostly EF-1α (21 leaves), enriched in pink conformations. Minor clusters included additional conformational states in light blue, ruby, and dark orange. Leaves in the tree were annotated with two inner rings: a thin inner ring indicating D81K-containing sequences, and a wider ring indicating WT-containing sequences. Sequences from D81A and D81F followed a pattern similar to D81K, therefore are not discussed separately. Most sequences belonged to canonical EF-Tu (gray), with smaller subsets classified as mitochondrial EF-Tu (blue), chlorophyll EF-Tu (yellow), and EF-1α (dark green).

A key observation is the significantly larger number of sequences retrieved using mutant queries: out of 1,318 total sequences, only 416 were from the WT query, while 902 were retrieved using mutant sequence as queries. Among EF-Tu entries, 379 sequences were shared between WT and D81K, 717 were unique to D81K, and only 9 were unique to WT. Taxonomic analysis reveals that the 379 shared sequences are predominantly from filamentous species, with 102 such entries (Figure 4F). In contrast, the 717 sequences uniquely recruited by D81K span a much broader ecological range, including anaerobic, chlorophototrophic, acidophilic, and thermophilic organisms (the outmost ring around the tree in Figure 4E, and Figure 4G). Many of these species thrive in extreme environments where translation is slower, allowing for greater EF-Tu variation.

### Paradox in MSA searching and structure prediction

Interestingly, although the retrieved MSA sequences in the D81K MSA largely retain the wild-type aspartate (D) at position 81 (LOGO plots in Figures 4E and S4), conformational clustering reveals a marked shift toward more restricted D-open structures compared to those associated with the WT query (blue conformations in Figure 4), as well as larger MSA sequence collections. This paradox highlights a novel effect: a single-point mutation introduced into the query sequence (D81K, D81A, or D81F), though not present in the aligned sequences themselves, significantly reshapes the sequence landscape sampled by AlphaFold. One possible explanation is that mutating a deeply conserved site like D81 perturbs its dominant co-evolutionary coupling network, thereby unmasking cryptic diversity in nearby regions that had been constrained by evolutionary pressure.^37–39^ In this view, the central mutation relaxes local constraints and shifts the attention landscape of AlphaFold, allowing it to detect alternative co-variation signals that are previously suppressed. This effect may have broader implications for protein evolution and ancestral reconstruction. It suggests that entrenched residues can obscure prior viable configurations, and that properly designed mutations in conserved positions may help recover functionally relevant conformational states lost during deep evolutionary time. Such an approach may offer a novel route for probing ancient protein diversity and refining models of molecular evolution.

## Discussion

In this study, we combined AlphaFold structure prediction, structure guided phylogenetics, and SURFS spectroscopy to investigate how mutations at the highly conserved D81 residue of EF-Tu alter conformational dynamics, sequence alignment behavior, and functional outputs.

Although D81 mutations did not impair GTP hydrolysis, as confirmed by fluorescence TLC measurements of mant-GDP levels, they significantly disrupted EF-Tu’s coordination of tRNA accommodation and ribosome movement. These results illustrate that GTP hydrolysis alone does not account for the functional impact of EF-Tu mutations. Instead, allosteric communication between the GTP binding site and the ribosome decoding center is critical and can be disrupted by a single allosteric mutation. These effects were detectable only using our SURFS platform, a force spectroscopy method developed to resolve ribosome conformational changes with sub-nucleotide resolution.

The underlying mechanism of these mutational effects was further explored through AlphaFold2 and AlphaFold3 simulations. While nucleotide binding remained stable in all models, the D81 mutants consistently showed weaker coordination of the magnesium ion and, more critically, impaired docking of the sarcin ricin loop (SRL) in the GTP bound state. Since SRL engagement is essential for ribosome mediated GTPase activation, this defect reflects a disruption in the long range structural communication required for signal propagation. These findings support a model in which D81 acts as a key coupling node that connects the GTP binding pocket to the decoding center. When disrupted, the resulting conformational decoupling alters ribosome function without affecting GTP hydrolysis.

Our findings further show that evolutionary conservation at D81 masks broader sequence and structural diversity in EF-Tu. Mutating this residue, even without propagating the mutation into the aligned sequences, weakens its dominance in the co-evolution network and allows cryptic structural features to emerge. The D81K mutation, in particular, expanded the accessible phylogenetic space to include anaerobic, acidophilic, chlorophototrophic, and thermophilic species, lineages in which EF-Tu may operate under relaxed constraints due to slower translation rates or altered cellular environments. This suggests that conserved residues can restrict our view of conformational and evolutionary diversity, and that probing beyond them opens a new window into both functional mechanisms and deep evolutionary history.

Our work suggests a new direction for probing ancient protein dynamics and recovering buried functional states. This is particularly valuable for multifunctional proteins like EF-Tu, which not only deliver aminoacyl-tRNAs to the ribosome but also moonlight in roles such as hibernation, cell-cell communication, and other regulatory processes beyond translation.^40–43^ By introducing targeted mutations at conserved residues and combining structure prediction with biophysical validation, we can possibly reconstruct lost conformational variants that may have existed in deep evolutionary time. The SURFS method provides the resolution needed to detect these subtle effects, which are undetectable to conventional enzymatic or structural assays. Future work will expand this strategy to other translation factors and universally conserved proteins.

### Limitations of the study

This study examines a limited set of EF-Tu mutants targeting a single conserved residue, which may not reflect the broader range of allosteric effects present across the protein. While SURFS provides high-resolution ribosome footprinting, it does not directly visualize structural rearrangements and depends on integration with predictive models. AlphaFold simulations, though informative, do not include full ribosomal context and require further experimental validation. The evolutionary analysis offers a testable hypothesis regarding coevolutionary constraints, but additional biochemical and structural studies are needed to confirm the proposed mechanisms.

## AUTHOR CONTRIBUTION

S.X. and Y.W. conceived and designed the experiments. Y.Z. conducted SURFS experiments. J.L.J. prepared EF-Tu mutants and GTPase assay. S.X. invented SURFS instrument. Y.W. conducted AlphaFold and phylogenetic analysis. All author wrote the paper together.

## ACKNOWLEDGEMENTS

This work was supported by US National Institutes of Health (5R01GM111452), and US National Science Foundation (BIO-2130427). J.L.J. was supported by the “HAMBP” (T32GM008280) predoctoral fellowship. Y. W. was supported by the Welch Foundation E-1721.

## DECLARATION OF COMPETING INTEREST

S.X. and Y.W. declare financial interest in UForce Biotechnology.

## RESOURCE AVAILABILITY

### Lead contact

Further information and requests for resources and reagents should be directed to and will be fulfilled by the lead contact, Yuhong Wang

### Materials availability

This study generated three EF-Tu mutants: D81A, D81F, and D81K. Plasmids are available upon request.

### code availability

The code is available at: https://github.com/ywang6000/EF-Tu-structure-clustering.

## METHODS DETAILS

### ET-Tu D81 mutants expression and purification

The D81 mutants were introduced into wild type EF-Tu pET-21b(+) plasmid using the Q5® Site Directed Mutagenesis Kit form New England Biolabs and mutagenesis primers ordered from IDT DNA. Cells from an overnight culture were incubated in LB containing 1 M sorbitol and 25 mM betaine until mid-log phase. Mid-log-phase cells (OD600 ∼0.6–0.8) were induced with 0.1 mM IPTG and harvested after 18 hours of growth at 30 °C. The cell pellets were collected by centrifugation at 4,000 ×*g* for 30 minutes at 4 °C. Cell pellets were resuspended in lysis buffer (50 mM Tris-HCl, 10 mM MgCl₂, 5 mM BME, 5 μM GDP, 15% glycerol, 60 mM NH₄Cl, pH 7.6) supplemented with a small amount of lysozyme crystals and incubated at −20 °C overnight. After thawing, the cells were lysed by sonication at 20% amplitude for a total of 2 minutes (10 seconds on, 20 seconds off). The lysate was clarified by centrifugation at 15,000 ×*g* for 30 minutes at 4°C. The cleared lysate was loaded onto a HisTrap FF 5 mL column pre-equilibrated with binding buffer (50 mM Tris-HCl, 10 mM MgCl_2_, 5 mM BME, 5 μM GDP, 300 mM NaCl, pH 7.6) and allowed to cycle for 1 hour. The column was washed with 10 column volumes of wash buffer (50 mM Tris-HCl, 10 mM MgCl_2_, 5 mM BME, 5 μM GDP, 300 mM NaCl, pH 7.6) to remove nonspecific proteins. The target proteins were eluted at approximately 250 mM imidazole using a 5–500 mM gradient on a GE ÄKTA Explorer 10 FPLC system. Eluted fractions were analyzed by SDS-PAGE to confirm purity. Wild type and D81F were subsequently buffer-exchanged into storage buffer (50 mM Tris-C, 10 mM MgCl2, 5 mM BME, 5 μM GDP, 100 mM NaCl, pH 7.6) while D81A and D81K underwent an additional purifying step by diluting the sample with Milli-Q water before loading on a HiTrap DEAE Sepharose FF column 1 mL and cycling overnight. The column was washed with 10 column volumes of wash buffer (50 mM Tris-HCl, 10 mM MgCl_2_, 5 mM BME, 5 μM GDP, pH 7.6). The target proteins were eluted at approximately 300 mM NaCl using a 0–500 mM gradient on a GE ÄKTA Explorer 10 FPLC system.

### Ribosome preparation

70S ribosomes were purified from *E. coli* MRE600 following previously described protocols.^44^ The cleared supernatant was layered onto a pre-cooled 1.1 M sucrose cushion in Buffer II (20 mM Tris-HCl, pH 7.6; 10 mM MgCl₂; 100 mM NH₄Cl; 6 mM β-mercaptoethanol (BME); 0.5 mM EDTA) at a 1:1 volume ratio. Ribosomes were pelleted by ultracentrifugation using a Beckman XL-80 ultracentrifuge with a Type 45 Ti Fixed-Angle rotor at 120,000 × *g*, 4°C, for 20 hours. The resulting pellet was gently rinsed with Buffer II and resuspended in the same buffer. The sucrose cushion ultracentrifugation step was repeated twice more, after which the ribosomes were resuspended in a minimal volume of Buffer II. The concentration of ribosomes was determined by measuring UV absorbance at 260 nm, using the conversion factor of 1 A260 unit = 23 pmol of 70S ribosomes. The purified ribosome solution was aliquoted, flash-frozen in liquid nitrogen, and stored at -80°C.

### Translational factors preparation

Ribosome factors (His-tagged IF1, IF2, IF3, IF4, EF-G, EF-Tu, methionyl- and total-tRNA synthetases) are prepared using standard protocols as previous described.^44^

### fMet-tRNA^fMet^ preparation

*E. coli* tRNAᶠᴹᵉᵗ was overexpressed in BL21 Star (DE3)pLysS cells from a pBluescript II SK (+) plasmid (Genscript) and purified on a Sepharose 4B column with a reverse ammonium sulfate gradient (methionine-accepting activity: 500 pmol/A260 unit). Recombinant 6×His-tagged *E. coli* methionyl-tRNA synthetase and methionyl-tRNAᶠᴹᵉᵗ formyltransferase were expressed and purified (Altuntop, M.E., Ly, C.T., and Wang, Y. *Biophys. J.* **99**, 3002-3009, 2010). The formyl donor, 10-formyltetrahydrofolate, was prepared as described before (Dubnoff, J. S. and Maitra, U. *Proc. Natl. Acad. Sci. USA* **68**, 318-323, 1971). Aminoacylation and formylation were performed as a one-pot reaction containing 100 mM Tris-HCl (pH 7.5), 4 mM ATP, 20 mM MgCl₂, 10 mM KCl, 150 μM L-methionine, 750 μM neutralized formyl donor, 7 mM BME, 20 μM tRNAᶠᴹᵉᵗ, 12 μM methionyl-tRNA synthetase, and 16 μM formyltransferase. The mixture was incubated at 37°C for 15 min, acidified to 0.3 M sodium acetate (pH 5.0), and purified by phenol-chloroform extraction, gel filtration (NAP-10, Cytiva), and ethanol precipitation. The final fMet-tRNAᶠᴹᵉᵗ was dissolved in 2 mM sodium acetate (pH 5.0) and stored at -80°C.

### Preparation of MFGPost Ribosome Complex

Three mixtures were prepared: the ribosome mix, TuWG mix, and FG mix. The ribosome mix contained 1 μM ribosomes, 1.5 μM each of initiation factors (IF1, IF2, IF3), 2 μM mRNA, 4 μM charged fMet-tRNAᶠᴹᵉᵗ, and 4 mM GTP. The TuWG mix consisted of 4 μM EF-Tu, 4 μM EF-G, 4 mM GTP, 4 mM 2-phosphoenolpyruvate (PEP), and 0.02 mg/mL pyruvate kinase. The FG mix included 100 mM Tris-HCl (pH 7.8), 20 mM MgCl₂, 1 mM EDTA, 4 mM ATP, 7 mM β-mercaptoethanol (BME), 10% (final concentration) *e coli* total tRNA synthetase, 50 A_260_ units/mL total tRNA, and 0.25 mM phenylalanine and glycine. Each mixture was preincubated separately at 37°C for 15 minutes and then combined in a 1:2:2 volume ratio at 37°C for 10 minutes. The resulting MFG-post ribosome complex was layered onto a 1.1 M sucrose cushion and purified by centrifugation at 200,000–400,000×*g* for 3 hours at 4°C using a Hitachi CS150FNX ultracentrifuge with an S140AT rotor.

### GTP hydrolysis assay with thin layer chromatograph (TLC)

Reaction mixtures contained 1 μM EF-Tu (wild type or mutant), 400 μM mant-GTP (Sigma-Aldrich), 1× TAM10 buffer (20 mM Tris-Cl pH 7.5, 10 mM MgAc, 30 mM NH_4_Cl, 70 mM KCl, 5 mM EDTA, 7 mM BME), 0.5 μM of ribosomes (cognate, near-cognate, or no mRNA) or no ribosomes, 0.5 μM charged Phe-tRNA^Phe^ or no tRNA. While in dim lighting, the reaction tubes were incubated at 37°C for 10 minutes. After incubation, 0.8 μL of each reaction mixture was carefully spotted onto a TLC plate using a micropipette. The plates were air-dried for approximately 10 minutes before being placed into TLC chambers. The mobile phase consisted of 3 mL 100% ethanol, 2 mL aqueous ammonia (saturated, ∼29%), 0.5 mL 1 M ammonium acetate, 4.5 mL H₂O, and 30 mg EDTA in H⁺-form. TLC plates were incubated in the chambers until the mobile phase traveled the full length of the plate (∼60 minutes). After development, the plates were removed, air-dried, and placed in a dark chamber for visualization using a UVP UVGL-25 Compact 4-watt UV Lamp with a spectral range of 254–365 nm. Images of the TLC plates were captured and analyzed using ImageJ. Regions of interest (ROIs) correspond to separated spots for mant-GTP and its hydrolysis product, mant-GDP, were defined. Fluorescence intensities were measured, and the percentage of hydrolyzed mant-GTP was calculated as the ratio of the intensity of mant-GDP to the sum of the intensities of mant-GDP and mant-GTP.

### SURFS assay

The sample well, with a bottom surface area of 4×3 mm^2^, was coated with biotin and incubated with 0.25 mg/mL streptavidin for 1 hour. The EF-Tu–GTP complex was formed by incubating 10 µM EF-Tu, 1 µM GTP, 1 µM PEP, 0.02 µM pyruvate kinase (PK), and 1 µM TAM10 at 37 °C for 10 minutes. Afterward, 7.4 µM K-tRNA was added and incubated at 37 °C for 10 minutes to yield the EF-Tu ternary complex with 5 µM EF-Tu. The ribosome complex formed through incubation at 37 °C for 30 minutes in a mixture containing 0.03 µM Post mRNA, 0.63 µM EF-Tu ternary complex, 1.5 mM GTP, 1.5 mM PEP, 0.15 mM ATP, and 0.02 mg/mL PK. The ribosome complex was immobilized on the surface via the 5′-end biotin on the mRNA by incubating for 1 hour. Then 1 μM probing DNA was added and incubated overnight to form duplexes with the uncovered mRNA. Streptavidin-coated magnetic beads (M280, Invitrogen) at 0.5 mg/mL were introduced into the sample well and incubated for 2 hours. Before the force measurements, the sample was magnetized for 2 minutes using a permanent magnet (∼0.5 T) and centrifuged (5427R, Eppendorf) at 820×*g* for 20 minutes to remove non-specific bound magnetic particles. Details of the SURFS experiments have been reported previously (Zeng, Y., Mao, Y., Chen, Y., Wang, Y. & Xu, S. *Chem. Commun.* **59**, 14855, 2023). Ultrasound at 1 MHz was applied onto a piezo disk to generate acoustic radiation force on the magnetic beads labeled on the DNA-mRNA duplexes. The amplitude of ultrasound was gradually increased; at each amplitude, the sample’s magnetic signal was measured by an atomic magnetometer.

When the force exceeds the dissociation force of the duplex, an abrupt decrease in magnetic signal will be observed. This is because after the magnetic beads dissociate from the surface, their magnetic dipoles will become random due to Brownian motion. Therefore, the dissociation force of the DNA-mRNA duplexes can be determined from the force spectra. Because the duplexes with different bp numbers show discrete dissociation forces under our experimental condition, the length of the duplexes can be precisely determined from the dissociation forces. In this work, each force spectrum was repeated at least three times to ensure reproducibility. The force resolution was typically 0.4 pN at 20 pN. The mRNA sequence was 5′-Bio-sg/A AAU UAA AUU AAA AAG GAA AUA AAA AUG UUU <u>GGA AAA AAG</u> UAC GUA AAU CUA CUG CUG AA-3′. The DNA sequence for probing the 3′ end of the mRNA was 5′-BiotinTEG\CT CAA GTG CAG TAG ATT TAC GTA CTT T\-3′; the DNA for the 5′ end was 5′-BioTEG\T_50_ TT TTT AAT TTA ATT T\-3′.

## Supplemental information

**Figure S1.**
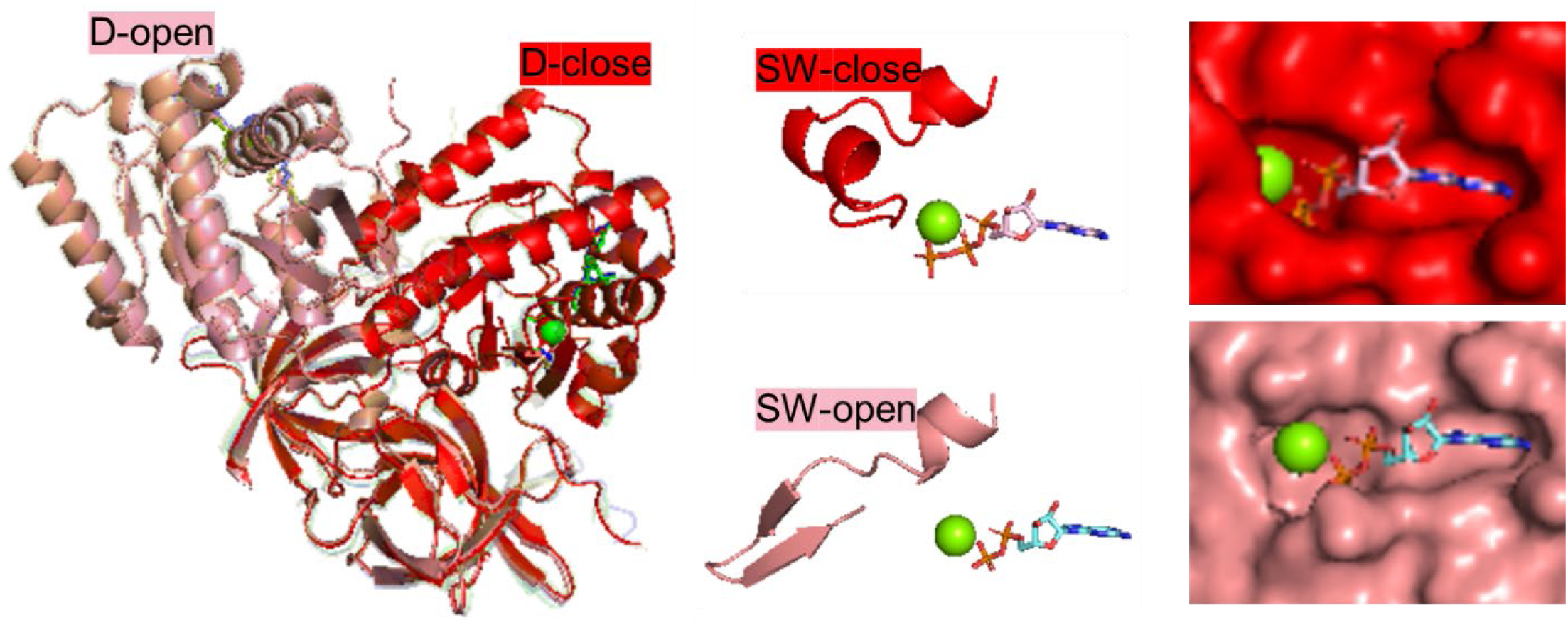
AlphaFold3 predicted EF-Tu complexes bound to GTP or GDP with Mg²⁺. The D and SW regions mark two conformationally flexible, allosterically connected sites. A 30-nucleotide sarcin–ricin loop (SRL) and a 22-nucleotide RNA were included in the modeling but are not shown for clarity.

**Figure S2.**
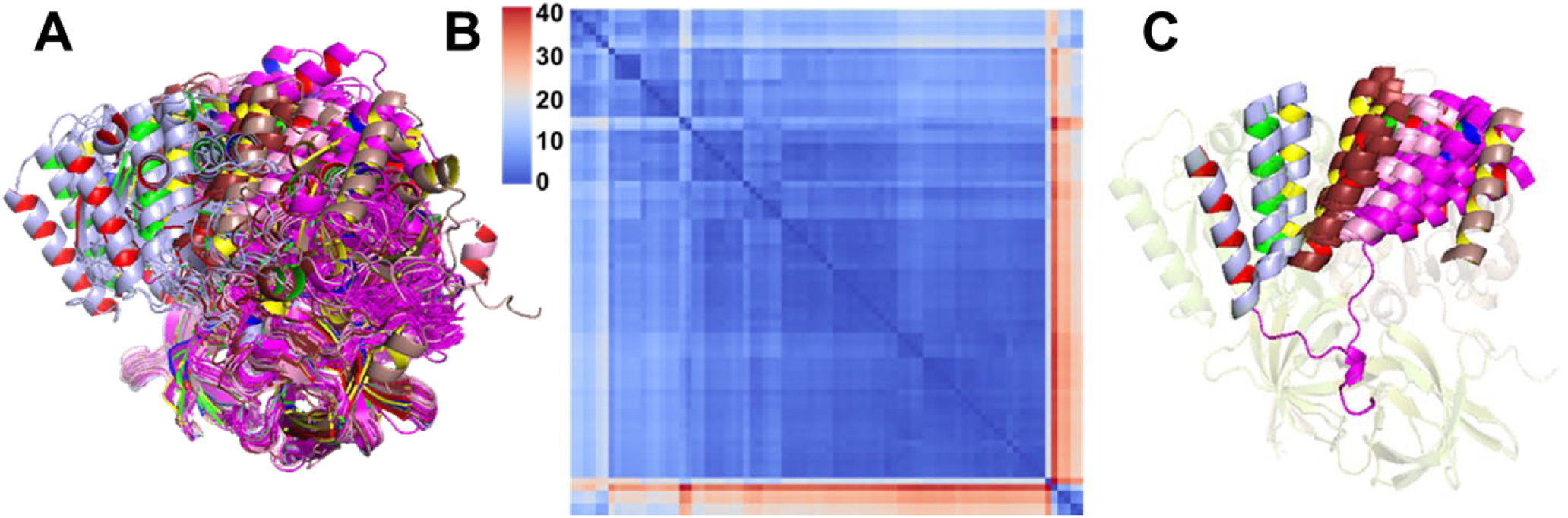
Structure clustering after AlphaFold2 prediction. (A) All high-confidence predicted structures aligned at domains II–III (residues 212–394). (B) RMSD matrix of the terminal helix in domain I (residues 183–199) for all structures shown in (A). (C) Representative conformations from each protein variant. Ribbon colors indicate conformation (outer) and protein identity (inner), respectively.

**Figure S3.**
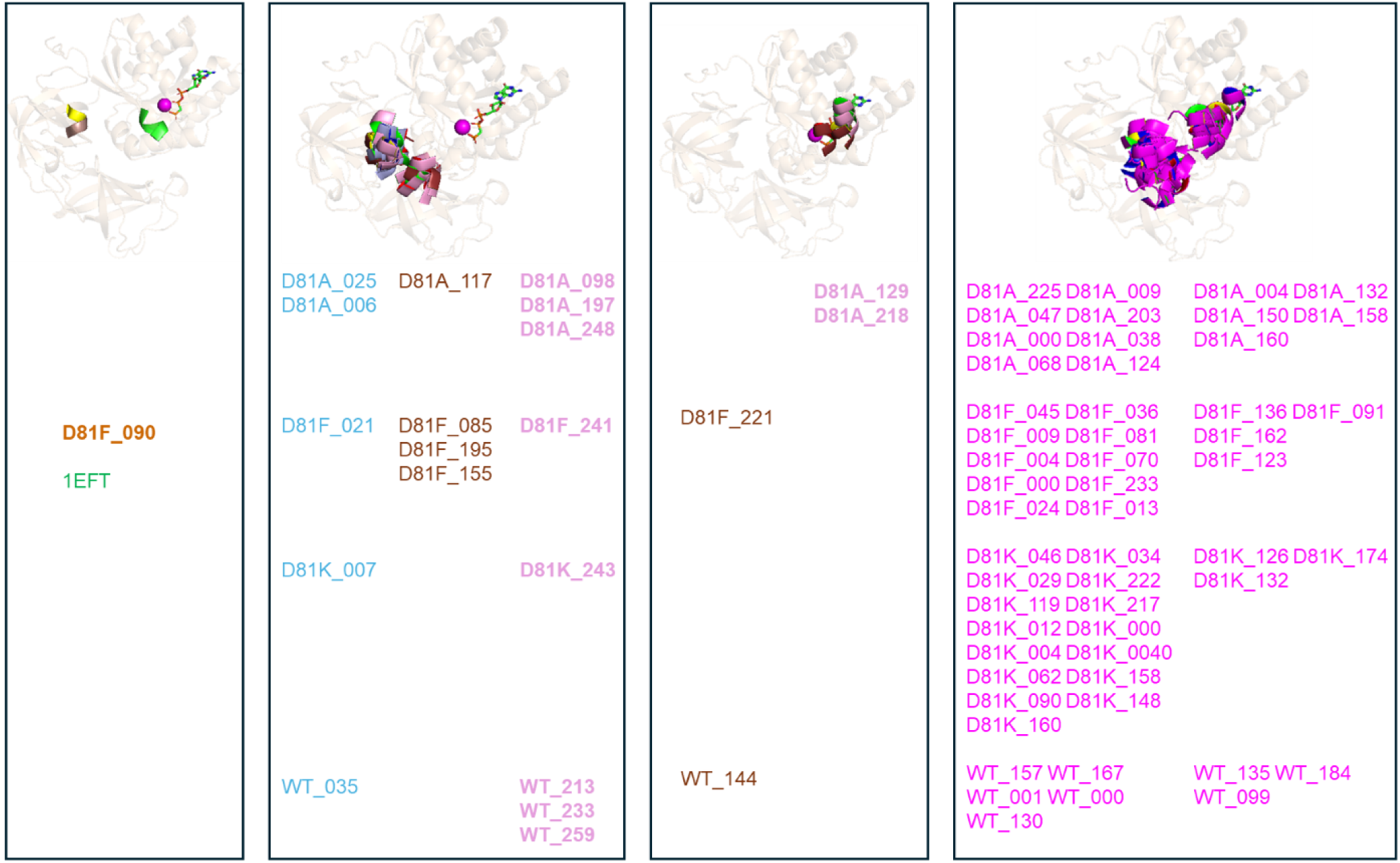
Conformations in the GTP-binding pockets of clustered structures from Figure S2. Color scheme matches the clusters shown in Figure S2. The background EF-Tu scaffold is based on the D-close form crystal structure (PDB: 1EFT). The first two panels showed SW-open forms. The third panel showed SW-close forms. The fourth panel showed both SW-open and close forms for magenta cluster (con3).

**Figure S4.**
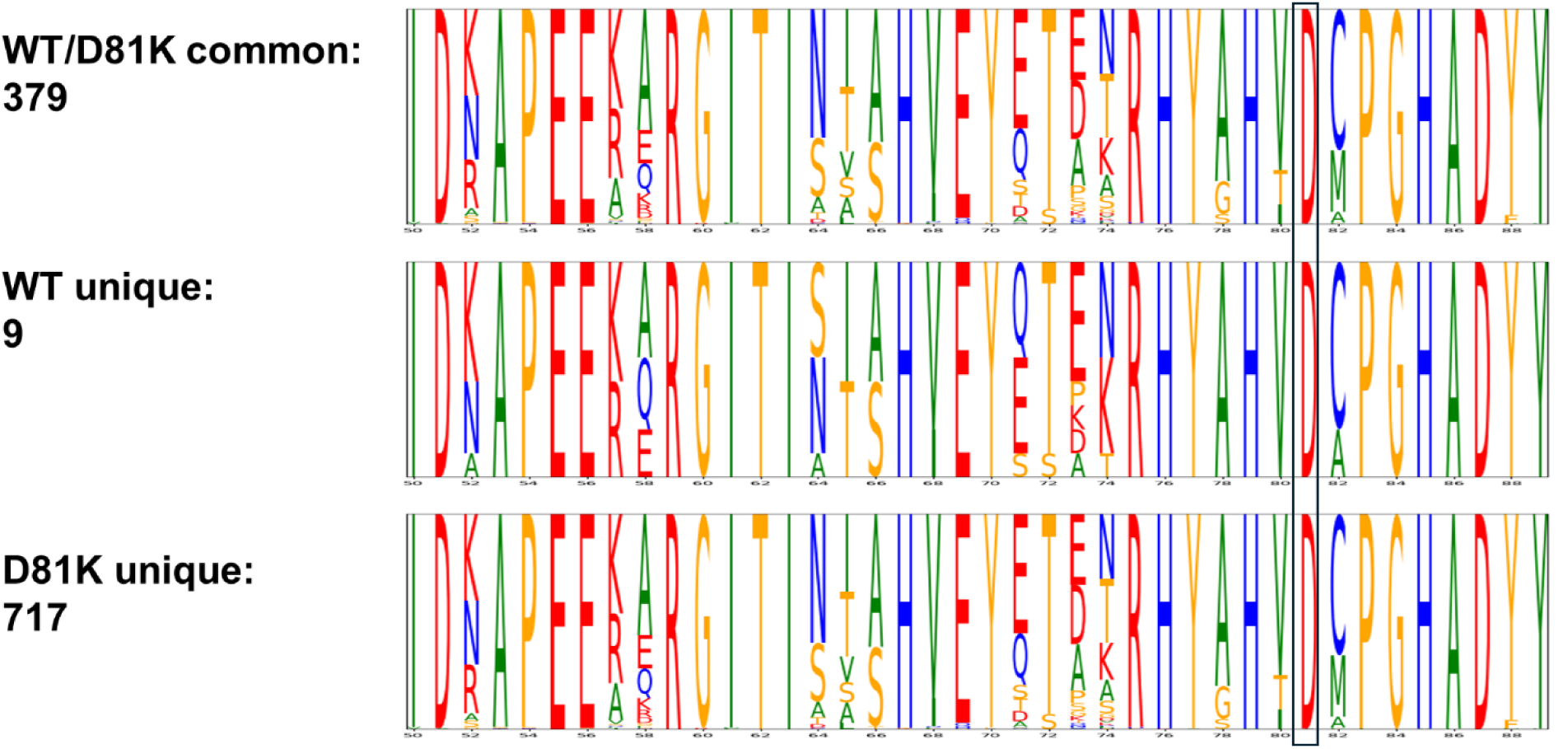
Position D81 is highlighted with a frame. Although these sequences span diverse taxonomic phyla, residue 81 remains conserved as aspartate (D), indicating that diverse MSA compositions can emerge from a mutated query even when the resulting MSA sequences retain the original residue at the mutated location.

**Table S1.**
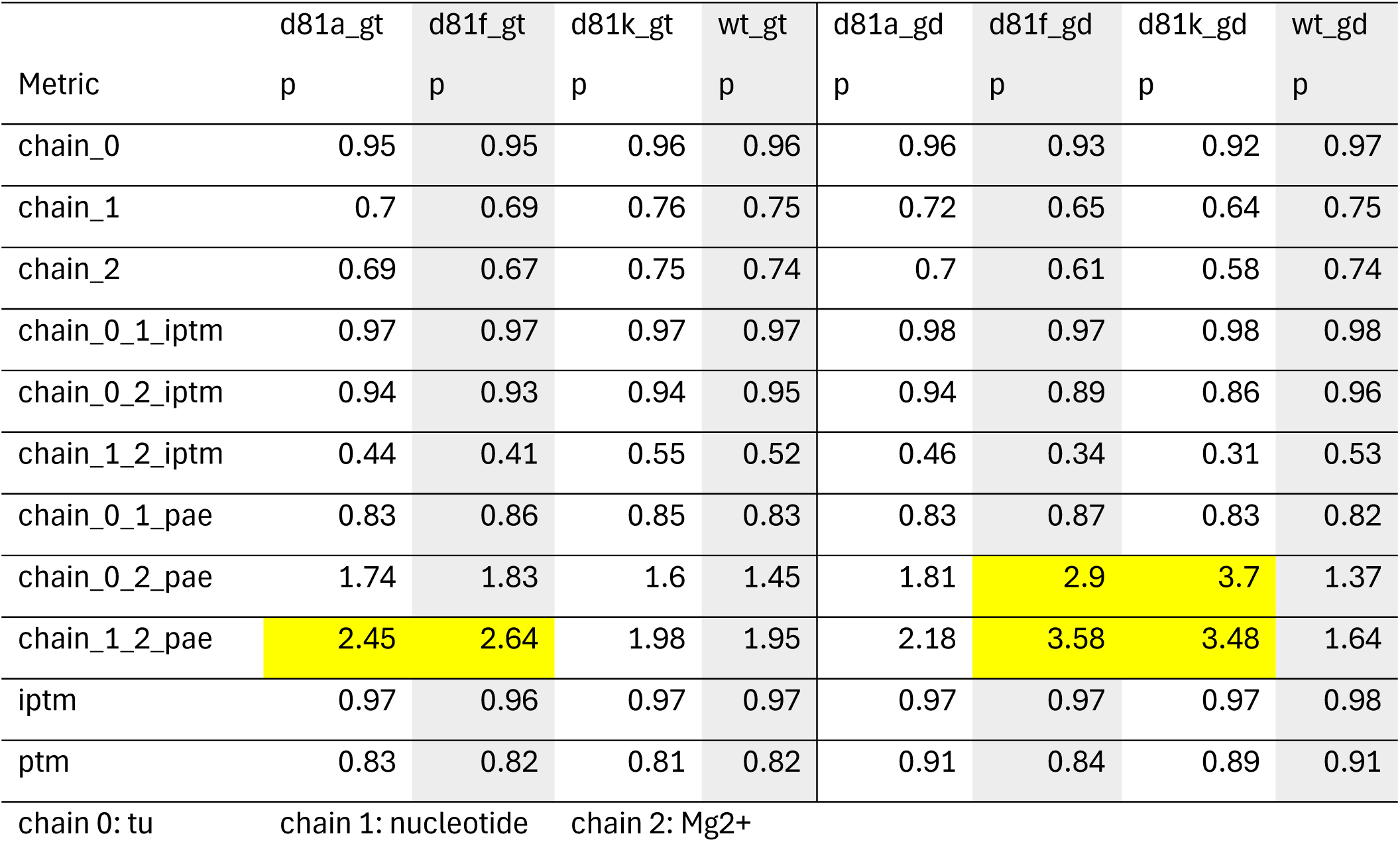
Folding confidence metrices for EF-Tu (chain 0) with nucleotide (chain 1) and Mg2+ (chain 2).

|  | d81a_gt | d81f_gt | d81k_gt | wt_gt | d81a_gd | d81f_gd | d81k_gd | wt_gd |
| --- | --- | --- | --- | --- | --- | --- | --- | --- |
| Metric | p | p | p | p | p | p | p | p |
| chain_0 | 0.95 | 0.95 | 0.96 | 0.96 | 0.96 | 0.93 | 0.92 | 0.97 |
| chain_1 | 0.7 | 0.69 | 0.76 | 0.75 | 0.72 | 0.65 | 0.64 | 0.75 |
| chain_2 | 0.69 | 0.67 | 0.75 | 0.74 | 0.7 | 0.61 | 0.58 | 0.74 |
| chain_0_1_iptm | 0.97 | 0.97 | 0.97 | 0.97 | 0.98 | 0.97 | 0.98 | 0.98 |
| chain_0_2_iptm | 0.94 | 0.93 | 0.94 | 0.95 | 0.94 | 0.89 | 0.86 | 0.96 |
| chain_1_2_iptm | 0.44 | 0.41 | 0.55 | 0.52 | 0.46 | 0.34 | 0.31 | 0.53 |
| chain_0_1_pae | 0.83 | 0.86 | 0.85 | 0.83 | 0.83 | 0.87 | 0.83 | 0.82 |
| chain_0_2_pae | 1.74 | 1.83 | 1.6 | 1.45 | 1.81 | 2.9 | 3.7 | 1.37 |
| chain_1_2_pae | 2.45 | 2.64 | 1.98 | 1.95 | 2.18 | 3.58 | 3.48 | 1.64 |
| iptm | 0.97 | 0.96 | 0.97 | 0.97 | 0.97 | 0.97 | 0.97 | 0.98 |
| ptm | 0.83 | 0.82 | 0.81 | 0.82 | 0.91 | 0.84 | 0.89 | 0.91 |
| chain 0: tu | chain 1: nucleotide |  | chain 2: Mg2+ |  |  |  |  |  |

**Table S2.** Folding confidence metrices for EF-Tu (chain 0) with nucleotide (chain 1), Mg2+ (chain 2), 22nt RNA (chain 3), and 30 nt SRL (chain 4).

| Metric | a81_gtp_sl | f81_gtp_sl | k81_gtp_sl | wt_gtp_sl | a81_gdp_sl | f81_gdp_sl | k81_gdp_sl | wt_gdp_sl |
| --- | --- | --- | --- | --- | --- | --- | --- | --- |
| chain_0 | 0.74 | 0.79 | 0.79 | 0.85 | 0.84 | 0.78 | 0.8 | 0.84 |
| chain_1 | 0.48 | 0.49 | 0.51 | 0.59 | 0.48 | 0.46 | 0.47 | 0.5 |
| chain_2 | 0.41 | 0.41 | 0.42 | 0.47 | 0.34 | 0.32 | 0.33 | 0.41 |
| chain_3 | 0.28 | 0.3 | 0.28 | 0.27 | 0.26 | 0.2 | 0.22 | 0.25 |
| chain_4 | 0.13 | 0.23 | 0.33 | 0.5 | 0.43 | 0.41 | 0.43 | 0.41 |
| chain_0_1_ip | 0.97 | 0.97 | 0.97 | 0.97 | 0.97 | 0.97 | 0.97 | 0.97 |
| chain_0_2_ip | 0.96 | 0.95 | 0.95 | 0.97 | 0.85 | 0.81 | 0.8 | 0.95 |
| chain_0_3_ip | 0.68 | 0.72 | 0.6 | 0.6 | 0.71 | 0.55 | 0.61 | 0.66 |
| chain_0_4_ip | 0.33 | 0.5 | 0.64 | 0.86 | 0.82 | 0.77 | 0.82 | 0.79 |
| chain_1_2_ip | 0.56 | 0.52 | 0.53 | 0.55 | 0.27 | 0.25 | 0.25 | 0.4 |
| chain_1_3_ip | 0.29 | 0.29 | 0.25 | 0.22 | 0.15 | 0.12 | 0.12 | 0.15 |
| chain_1_4_ip | 0.09 | 0.19 | 0.29 | 0.61 | 0.51 | 0.52 | 0.53 | 0.48 |
| chain_2_3_ip | 0.06 | 0.06 | 0.05 | 0.05 | 0.02 | 0.02 | 0.02 | 0.03 |
| chain_2_4_ip | 0.04 | 0.09 | 0.16 | 0.32 | 0.21 | 0.21 | 0.23 | 0.24 |
| chain_3_4_ip | 0.07 | 0.13 | 0.2 | 0.21 | 0.16 | 0.12 | 0.14 | 0.15 |
| chain_0_1_pa | 0.88 | 0.95 | 0.9 | 0.88 | 0.94 | 0.92 | 0.91 | 0.95 |
| chain_0_2_pa | 1.31 | 1.56 | 1.52 | 1.2 | 3.65 | 4.62 | 4.98 | 1.55 |
| chain_0_3_pa | 4.54 | 4.11 | 6.4 | 6.87 | 3.71 | 7.71 | 4.83 | 5.11 |
| chain_0_4_pa | 14.17 | 10.43 | 6 | 1.83 | 2.16 | 3.1 | 2.42 | 2.91 |
| chain_1_2_pa | 1.82 | 2 | 2.01 | 1.7 | 3.98 | 4.54 | 4.88 | 2.18 |
| chain_1_3_pa | 5.65 | 5.41 | 7.94 | 7.59 | 7.47 | 11.73 | 9.22 | 8.6 |
| chain_1_4_pa | 15.77 | 13.78 | 9.71 | 1.98 | 2.53 | 3.24 | 2.7 | 2.77 |
| chain_3_4_pa | 16.16 | 13.38 | 10.81 | 11.25 | 9.62 | 14.17 | 12.34 | 10.88 |
| iptm | 0.77 | 0.76 | 0.78 | 0.84 | 0.82 | 0.77 | 0.8 | 0.8 |
| ptm | 0.83 | 0.81 | 0.83 | 0.84 | 0.87 | 0.83 | 0.84 | 0.85 |
| chain0: Tu | 1: nucleoti | 2: Mg2+ | 3: RNA 22r | 4: SRL 30 nt |  |  |  |  |

**Table S3.**
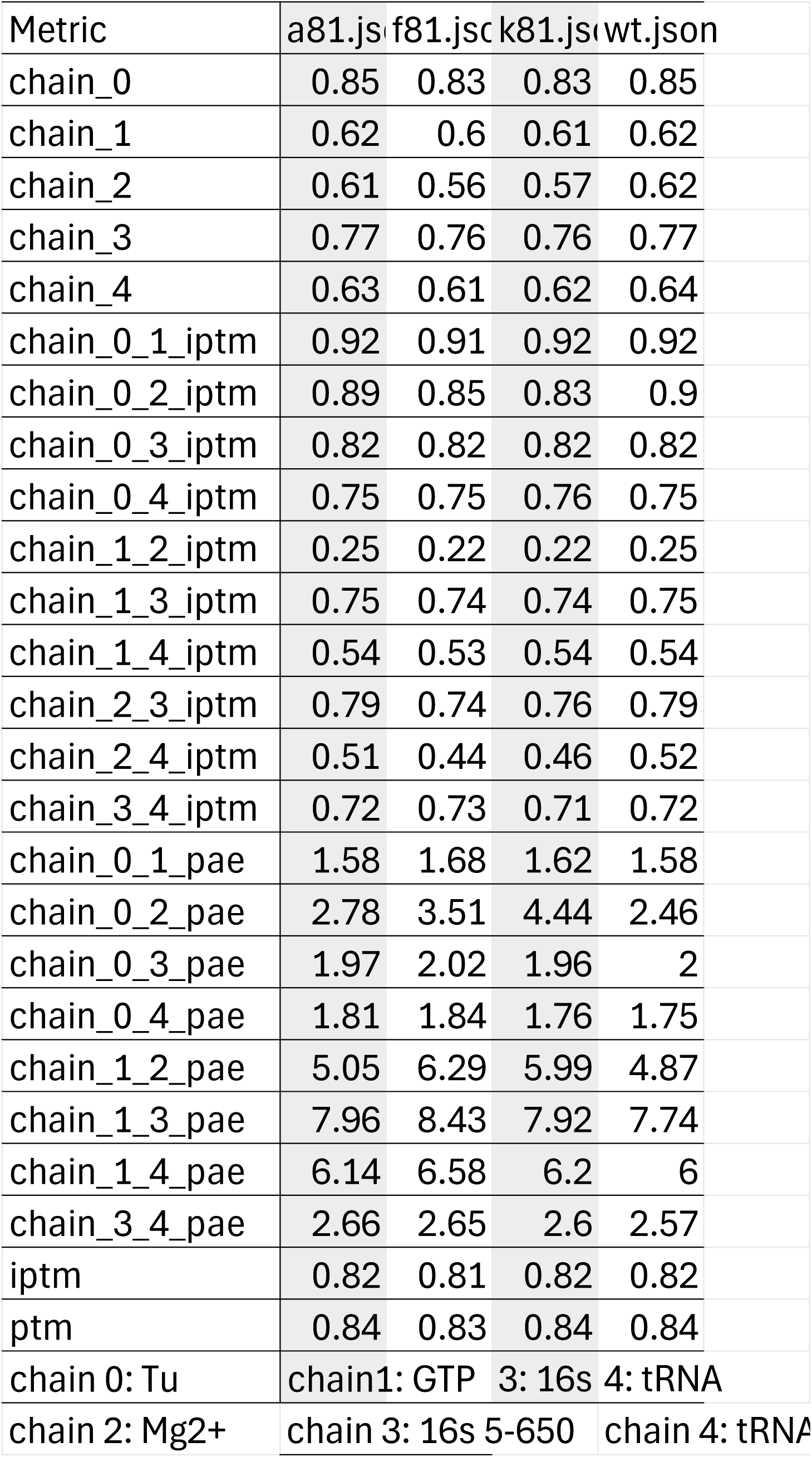
Folding confidence metrices for EF-Tu (chain 0) with GTP (chain 1), Mg2+ (chain 2), partial 16S RNA (chain 3), and tRNA (chain 4).

## Notes

https://github.com/ywang6000/EF-Tu-structure-clustering

